# Integrated pharmacological and structural profiling reveals CB1 residue interaction patterns associated with synthetic cannabinoid receptor agonist efficacy

**DOI:** 10.64898/2026.09.08.748791

**Authors:** Hyeokjun Kwon, Chul Kyu Lee, Jae-Hwan Kwak, Jaesuk Yun

## Abstract

**Background and Purpose:** Among new psychoactive substances, synthetic cannabinoid receptor agonists (SCRAs) encompass considerable structural diversity and show wide variation in cannabinoid receptor 1 (CB1) potency and efficacy, but the molecular and receptor-interaction characteristics underlying these differences remain unclear. We assessed the pharmacological profiles of 16 SCRAs and examined structural features associated with differences in CB1 potency and efficacy.

**Experimental Approach:** CB1 agonist activity was measured in a CB1-Ga15-based Ca^2+^ assay, and cataleptic effects were assessed in mice. Ligand-receptor interactions were characterized by molecular docking to an active-state CB1 structure. Principal component analysis (PCA) was applied to residue-level interaction profiles, and the resulting scores were tested for associations with CB1 potency and efficacy.

**Key Results:** Potency and efficacy varied widely across cellular and animal assays, with several SCRAs exhibiting different pharmacological responses across the two experimental systems. PCA identified an interaction pattern whose PC1 scores were significantly associated with *E*_max_ but not EC_50_, with contributions from activation-related CB1 residues including the PHE200-TRP356 toggle switch. Molecular descriptor analysis likewise identified structural features significantly associated with *in vitro* and *in vivo E*_max_.

**Conclusion and Implications:** Our findings suggest that CB1 efficacy is associated with distinct receptor-interaction patterns among structurally diverse SCRAs. Combining functional pharmacology with receptor-level interaction analysis provides structural insight into SCRA efficacy and may help prioritize emerging compounds for further pharmacological and behavioral evaluation.

**Bullet Point Summary:** *What is already known:*

- SCRAs act at CB1 receptors and vary widely in potency and efficacy.
- Structurally similar SCRAs can produce markedly different pharmacological effects.

*What this study adds:*

- PC1 derived from residue-interaction profiles was significantly associated with *in vitro E*_max_ in 11 SCRAs.
- Selected molecular descriptors and PC1 were associated with *E*_max_, but not EC_50_.

*Clinical significance:*

- Efficacy-linked interaction patterns may help prioritize emerging SCRAs for pharmacological and behavioral evaluation.

## Introduction

New psychoactive substances (NPS) include analogues of controlled drugs and newly developed chemicals designed to reproduce the psychoactive effects of illicit substances or pharmaceutical products [1–3]. New compounds continually emerge in illicit markets, and their chemical diversity presents substantial challenges for public health, forensic toxicology, clinical management, and drug control worldwide [3–5].

Synthetic cannabinoid receptor agonists (SCRAs) from a prominent NPS category displaying substantial structural diversity; more than 290 distinct compounds have been documented by the United Nations Office on Drugs and Crime [6–8]. Their psychoactive activity is mediated primarily through cannabinoid receptor 1 (CB1), which belongs to the G protein-coupled receptor family and is broadly distributed throughout the central nervous system [9]. Unlike Δ-tetrahydrocannabinol (Δ_9_-THC), which accounts for much of the psychoactive activity of cannabis and exhibits partial agonism at CB1, many SCRAs acts as high-efficacy full agonists and produce stronger pharmacological effects [10,11]. Their use has been associated with dependence, psychosis, seizures, organ toxicity, and death [12]. In animal models, SCRAs such as JWH-018 elicit the cannabinoid tetrad, characterized by hypothermia, analgesia, reduced locomotor activity, and catalepsy [13]. However, many SCRAs remain pharmacologically uncharacterized, and structurally similar compounds can show markedly different CB1 activity [14].

Subtle changes in SCRA molecular architecture may therefore alter CB1 pharmacology, supporting systematic structure-activity relationship analysis. Chemoinformatic approaches, including molecular fingerprint similarity analysis and hierarchical clustering, can classify structurally related NPS and predict their chemical and pharmacological characteristics [15–17].

Active-state CB1 structures show that ligand binding induces conformational changes, with PHE200 and TRP356 forming a toggle switch involved in activation, and TRP279 and other binding-pocket residues contributing to ligand binding and stabilization [18]. Molecular docking can therefore compare how structurally diverse SCRAs engage functionally relevant CB1 residues. However, the relationship between these interactions and SCRA potency or efficacy remains poorly understood.

Functional assays are needed to determine whether structural differences affect CB1 activation. CB1 couples mainly to G_i/o_ proteins [19]; coupling it to the promiscuous G protein Ga15 allows activation to be measured through intracellular Ca^2+^ mobilization and yields EC_50_ and *E*_max_values [20]. Complementary *in vivo* assessment is important because receptor-level activity may not directly translate into behavioral effects. Catalepsy, one of the cannabinoid tetrad effects, provides a measure of CB1-mediated activity in animals [13].

We selected 16 SCRAs for their regulatory relevance across multiple jurisdictions [6–8,21–26] and evaluated them using *in vitro* CB1 functional profiling and *in vivo* catalepsy testing. We compared the *in vitro* and *in vivo* responses. Ligand–residue interaction profiles obtained by molecular docking were analyzed by hierarchical clustering and PCA. The primary aim was to identify CB1 residue interaction patterns associated with differences in SCRA efficacy.

## Methods

### Drugs

The National Institute of Food and Drug Safety Evaluation (NIFDS), Ministry of Food and Drug Safety, supplied 16 structurally diverse SCRAs: JWH-018 (reference compound) [naphthalen-1-yl-(1-pentylindol-3-yl)methanone], JWH-210 [(4-ethylnaphthalen-1-yl)-(1-pentylindol-3-yl)methanone], MDA-19 [N’-[(3Z)-1-Hexyl-2-oxo-1,2-dihydro-3H-indol-3-ylidene]benzohydrazide], 5F-MDA-19 [N’-(1-(5-fluoropentyl)-2-oxoindolin-3-ylidene)benzohydrazide], AKB-48 [N-(1-adamantyl)-1-pentyl-1H-indazole-3-carboxamide], FUBIMINA [(1-(5-fluoropentyl)-1H-indazole-3-yl)(naphthalen-1-yl)methanone], 5F-PCN [1-(5-fluoropentyl)-*N*-naphthalen-1-ylpyrrolo[3,2-c]pyridine-3-carboxamide], 5F-MDMB-P7AICA [Methyl(2S)-2-[[1-(5-fluoropentyl)pyrrolo[2,3-b]pyridine-3-carbonyl]amino]-3,3-dimethylbutanoate], CUMYL-5F-P7AICA [1-(5-fluoropentyl)-*N*-(2-phenylpropan-2-yl)pyrrolo[2,3-b]pyridine-3-carboxamide], CUMYL-4CN-B7AICA [1-(4-Cyanobutyl)-N-(2-phenylpropan-2-yl)pyrrolo[2,3-b]pyridine-3-carboxamide], CB-13 [Naphthalen-1-yl-(4-pentyloxynaphthalen-1-yl)methanone], 5F-3,5-AB-PFUPPYCA [*N*-[(2*S*)-1-amino-3-methyl-1-oxobutan-2-yl]-1-(5-fluoropentyl)-3-(4-fluorophenyl)pyrazole-5-carboxamide], LY-2183240 [*N*,*N*-dimethyl-5-[(4-phenylphenyl)methyl]tetrazole-1-carboxamide], EG-018 [naphthalen-1-yl-(9-pentyl-9H-carbazol-3-yl)methanone], CUMYL-PEGACLONE [5-pentyl-2-(2-phenylpropan-2-yl)pyrido[4,3-b]indol-1-one], and 5F-CUMYL-PEGACLONE [5-(5-fluoropentyl)-2-(2-phenylpropan-2-yl)pyrido[4,3-b]indol-1-one]. Lipomed (Arlesheim, Switzerland) supplied Δ^9^-tetrahydrocannabinol [(6aR,10aR)-6,6,9-trimethyl-3-pentyl-6a,7,8,10a-tetrahydrobenzo[c]chromen-1-ol] (△_9_-THC). Toprak Mahsulleri Ofisi (Ankara, Turkiye) supplied cocaine hydrochloride [Methyl (1R,2R,3S,5S)-3-benzoyloxy-8-methyl-8-azabicyclo[3.2.1]octane-2-carboxylate;hydrochloride] which served as a control for assay specificity. Stock solutions in distilled water or dimethyl sulfoxide (DMSO) were maintained at -80 □ and diluted with assay buffer prior to experiments.

### Animals

All procedures involving animals were conducted according to the Guide for the Care and Use of Laboratory Animals and approved by the Animal Care Committee of Chungbuk National University (Cheongju, Republic of Korea; approval CBNUA-2027-22-02). Male C57BL/6N mice were supplied by Daehan Biolink (Eumsung, Republic of Korea). Four animals were housed in each cage and used at 8–9 weeks, with room conditions set to 23 ± 2 °C, 50 ± 5% relative humidity, and a 12-h light/dark cycle. The light phase ran from 7 AM, while food and water remained freely available.

### Hierarchical clustering and fingerprint similarity analysis

Molecular fingerprint analysis was used to evaluate structural similarity among the synthetic cannabinoids. Canonical SMILES strings were obtained and converted into molecular structures using the RDKit cheminformatics toolkit [27]. Molecular fingerprints were generated using the RDKit fingerprint algorithm, which encodes substructural features as binary vectors. Pairwise similarity was calculated using the Tanimoto coefficient, representing shared features relative to the total features of two fingerprints. The similarity matrix was converted to a distance matrix (1 - similarity) and analyzed by hierarchical clustering using Ward linkage and Euclidean distance. Results were visualized as a dendrogram combined with a heatmap to identify structurally related groups.

### Cell culture

CB1-Ga15 stably expressing Chem-1 cells were obtained from Eurofins DiscoverX Products LLC (Fremont, CA, USA). Cells were grown at 37 □ under 5% CO_2_ in high-glucose DMEM (Cytiva, MA, USA) containing 10% fetal bovine serum (FBS, Cytiva, MA, USA), 1X non-essential amino acids (Cytiva, MA, USA), 1X HEPES (Merck KGaA, Darmstadt, Germany), and 1% penicillin-streptomycin (Gibco, NY, USA). Upon reaching confluence, cells were maintained in the G418-supplemented medium (250 pg/mL; InvivoGen, CA, USA). Cells at passage 3 were used for all experiments.

### Ca^2+^ assay

Chem-1 cells stably expressing CB1-Ga15 were seeded in poly-L-lysine-coated 96-well plates at 3 × 10^4^ cells/well and loaded with Fluo-8 AM (AAT Bioquest, CA, USA). Cells were incubated for 1 h at 37 □ with 5% CO_2_, washed three times with Hank’s balanced salt solution (HBSS; Sigma-Aldrich, MO, USA), and supplemented with 60 μ L of HBSS. Following a 60 s baseline recording, 30 pL of vehicle or drug (0.001 nM-100 pM) was applied. Fluorescence was then monitored for 60 s with a FlexStation 3 Multi-mode Microplate Reader (Molecular Devices, CA, USA) using excitation/emission wavelengths of 495/516 nm. Relative fluorescence changes (% ΔF/F_0_) were calculated by subtracting normalized baseline fluorescence from each post-treatment value.

### Catalepsy test

Cataleptic response was evaluated by the bar test. Each mouse placed both forelimbs on a 0.5 cm-diameter horizontal bar positioned 4.0 cm from the chamber floor. Posture duration was recorded with a 300 s cut-off. Measurements were performed 90 min after administration of vehicle or drug at 0.1, 0.2, 0.5, 1.0, 2.0, or 4.0 mg/kg, with three mice used per group. A vehicle consisting of saline, 5% DMSO, and 5% Tween 80 was used for drug administration.

### Molecular docking

Molecular docking was performed to characterize how the 16 SCRAs interact with and bind to CB1. The active-state cryo-EM structure of CB1 coupled to heterotrimeric G_i_ protein (PDB ID: 6N4B) served as a template for receptor modeling. The receptor was prepared using Schrödinger Maestro (Schrödinger, Inc., NY, USA), and the 16 SCRAs were prepared as ligands. Docking was performed using Glide extra-precision mode, with the receptor grid centered on the orthosteric ligand-binding pocket of CB1 as previously described [18]. Multiple poses were generated for each SCRA and ranked by their Glide docking scores. A representative pose was selected for each compound based on the docking results and compatibility with the CB1 ligand-binding pocket. Protein-ligand interactions in the selected poses were then analyzed to characterize residue-level interaction patterns.

### Principal component analysis

Residue-level interaction profiles obtained from molecular docking were subjected to PCA. Eleven SCRAs with reliably determined *in vitro* EC_50_ and *E*_max_ values were included. Seven residues within the CB1 ligand-binding pocket, HIS178, ARG182, PHE189, PHE200, PHE268, TRP279, and TRP356, were selected based on their reported roles in ligand binding and receptor activation. Interactions between each SCRA and these residues were classified from the representative docking poses as no interaction, other interaction, or tt-tt interaction to generate residue-level interaction fingerprints. PCA was performed using these fingerprints to identify major patterns of variation in CB1-SCRA interactions. The resulting principal component scores were compared with the experimentally determined *in vitro* EC_50_ and *E*_max_ values.

### Data and statistical analyses

Mean values are accompanied by their standard errors (SEM). Responses from both functional assays were expressed relative to the maximum JWH-018 responses, set at 100%. Dose-response curves were fitted by four-parameter logistic nonlinear regression to determine EC_50_ or ED_50_ and *E*_max_ values, representing potency and efficacy, respectively. Curve fits with R^2^ < 0.7 were considered insufficiently reliable, and the corresponding parameters were designated not determined (ND). GraphPad Prism 5 (version 5.02; GraphPad Software, CA, USA) was used for curve fitting and correlation analyses, and SigmaPlot 12.5 (Systat Software, CA, USA) was used for descriptive statistics and statistical power analyses.

The Shapiro-Wilk test assessed whether data followed a normal distribution. Pearson’s correlation evaluated continuous variables satisfying this assumption, whereas Spearman’s rank correlation evaluated those deviating from it. Spearman’s correlation further examined relationships between molecular descriptors and pharmacological parameters and between principal component scores (PC1, PC2, and PC3) and *in vitro* EC_50_ and *E*_max_. Benjamini-Hochberg adjustment was applied to Spearman correlation *P*-values for false discovery rate (FDR) correction, with *q* < 0.05 indicating significance. All analyses employed two-sided testing unless stated otherwise.

## Results

### Structural classification of the 16 SCRAs

To examine structural determinants of pharmacological effects, the 16 SCRAs were classified by molecular architecture (Figs. 1 and 2). Using JWH-018 as the reference backbone, each compound was divided into a core, linker (L1), head (R1), tail (R2), and additional residues (R3 and R4). Compounds containing indole, indazole, benzimidazole, 4-azaindole, and 7-azaindole cores were assigned to Classes 1–5, respectively, whereas those containing naphthalene, pyrazole, tetrazole, carbazole, and pyridoindole cores were assigned to Classes 6–10. Compounds were then subdivided by linker, head, tail, and residue groups. Linkers were classified as La– Ld, with La’ indicating an alternative attachment position; head groups as Ha–Hh, including positional variants Ha’ and Hf’; tails as Ta–Te; and additional residues as Ra–Rd.

**Fig. 1.**
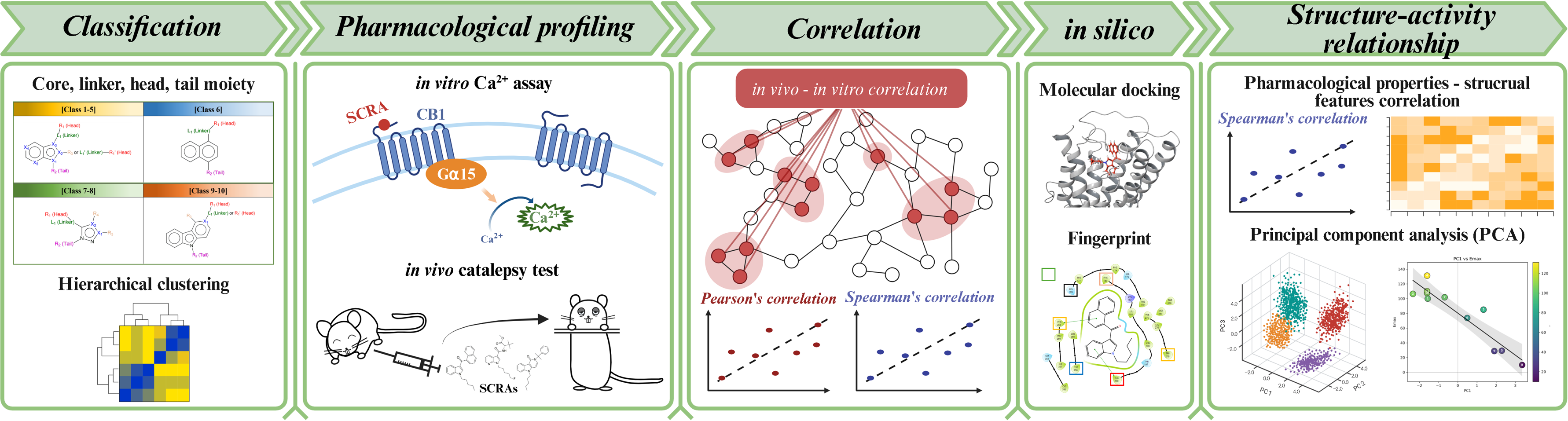
Molecular structures of SCRAs. The structures of 16 SCRAs are shown.

**Fig. 2.**
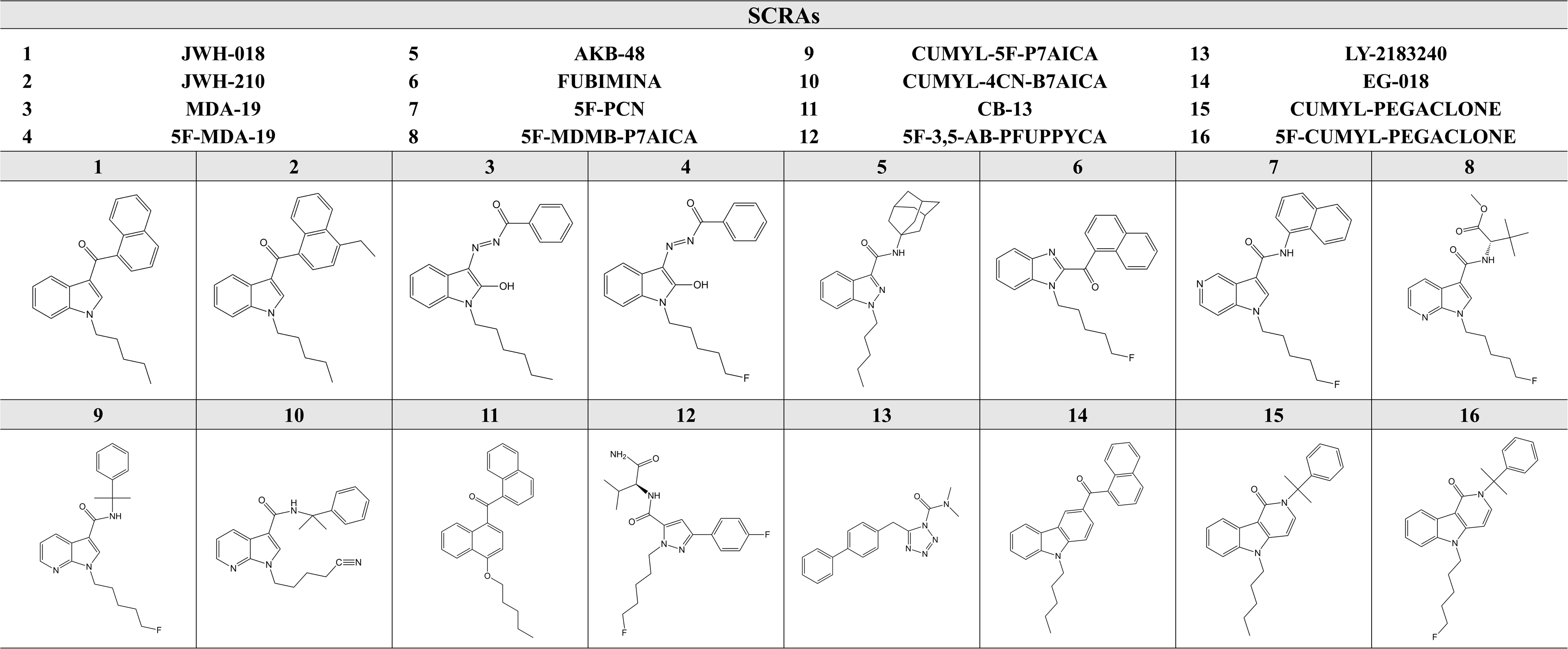
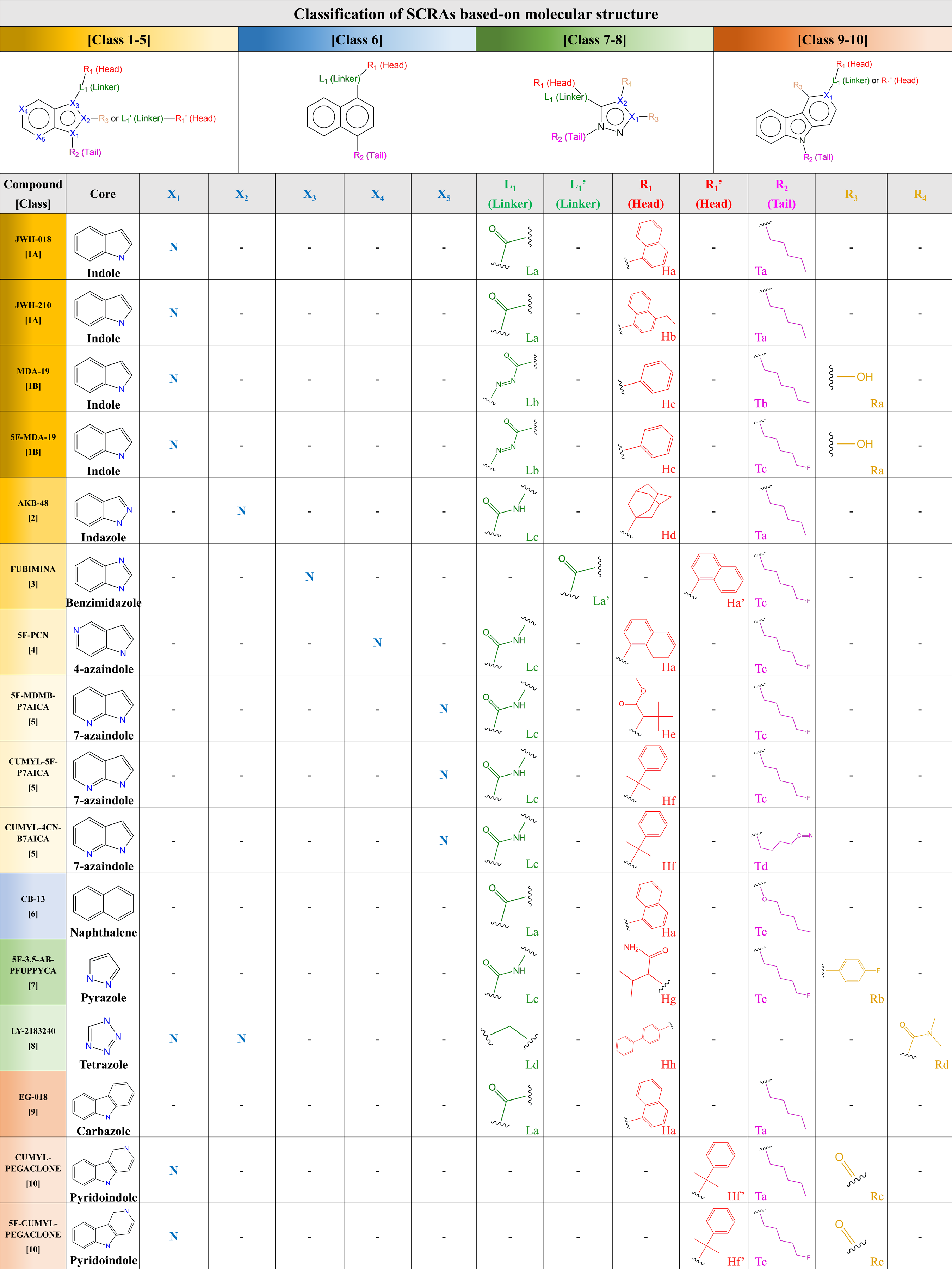
Classification of SCRAs based on molecular structure. SCRAs were classified according to their core scaffold and substituent groups, including linker (L1 and L1’), head (R1 and R1’), tail (R2), and additional residues (R3 and R4). Based on the core structure, compounds were grouped into Classes 1–10, including indole (Class 1), indazole (Class 2), benzimidazole (Class 3), 4-azaindole (Class 4), 7-azaindole (Class 5), naphthalene (Class 6), pyrazole (Class 7), tetrazole (Class 8), carbazole (Class 9), and pyridoindole (Class 10). Further subclassification was performed based on variations in linker, head, tail, and residue moieties, as indicated by the assigned structural codes (e.g., La-Ld, Ha-Hh, Ta-Tf, Ra-Rd).

JWH-018 and JWH-210 formed Class 1A, whereas MDA-19 and 5F-MDA-19 formed Class 1B, sharing an indole core but differing in linker structure. AKB-48, FUBIMINA, and 5F-PCN were assigned to Classes 2, 3, and 4, respectively. 5F-MDMB-P7AICA, CUMYL-5F-P7AICA, and CUMYL-4CN-B7AICA formed Class 5. CB-13, 5F-3,5-AB-PFUPPYCA, LY-2183240, and EG-018 were assigned to Classes 6–9, respectively, and CUMYL-PEGACLONE and 5F-CUMYL-PEGACLONE to Class 10. This structural classification provided a systematic framework for describing structural diversity among the tested SCRAs.

### Structural clustering supports the classification scheme

Hierarchical clustering based on molecular fingerprint similarity revealed distinct structural groupings among the SCRAs (Fig. 3). Compounds sharing core scaffolds, such as indole- and indazole-based SCRAs, clustered together. The 7-azaindole derivatives formed a compact cluster, whereas compounds differing in head or tail groups were distributed across multiple clusters. Some compounds with similar cores appeared in different clusters, corresponding to differences in their linker or residue groups. Overall, the clustering was largely consistent with the predefined classification based on core, linker, head, tail, and residue features.

**Fig. 3.**
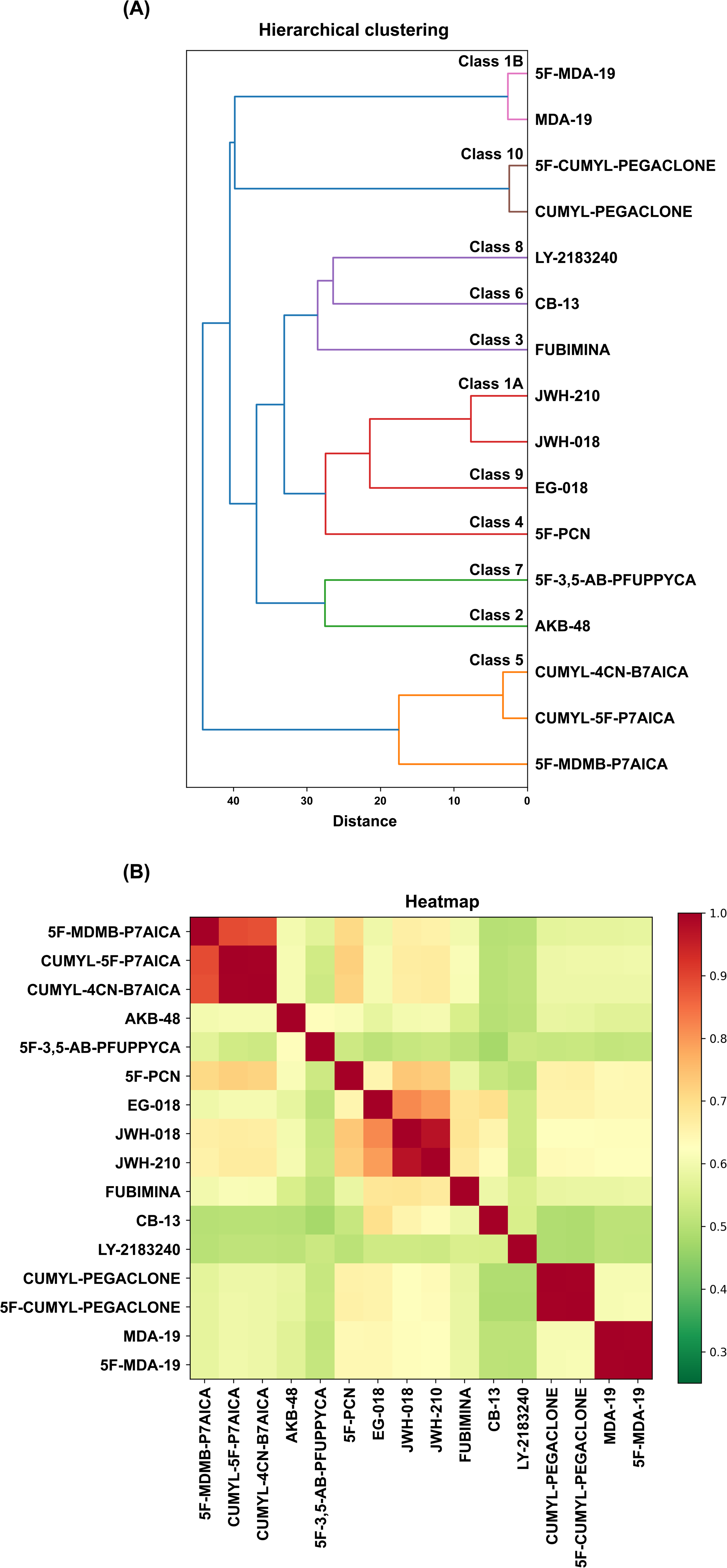
Structural similarity analysis of SCRAs using hierarchical clustering and heatmap visualization. A) Hierarchical clustering dendrogram of 16 SCRAs based on molecular fingerprint similarity. Canonical SMILES were converted into molecular fingerprints using the RDkit algorithm, and pairwise similarity was calculated using the Tanimoto coefficient. The resulting similarity matrix was transformed into a distance matrix (1-similarity), and clustering was performed using the Ward linkage method. B) Heatmap representation of pairwise structural distance among SCRAs. The color scale indicates the degree of structural similarity between compounds.

### SCRAs show wide differences in CB1 potency and efficacy *in vitro*

To validate the *in vitro* assay, the full CB1 agonist JWH-018, the partial agonist A^9^-THC, and cocaine, a dopamine transporter inhibitor used as a negative control, were tested (Figs. 4A and S1). JWH-018 elicited concentration-dependent intracellular Ca^2+^ signaling, while A9-THC produced a smaller maximal response, consistent with partial CB1 agonism. Cocaine produced no CB1-mediated Ca^2+^ response, indicating assay specificity (Fig. 4A).

**Fig. 4.**
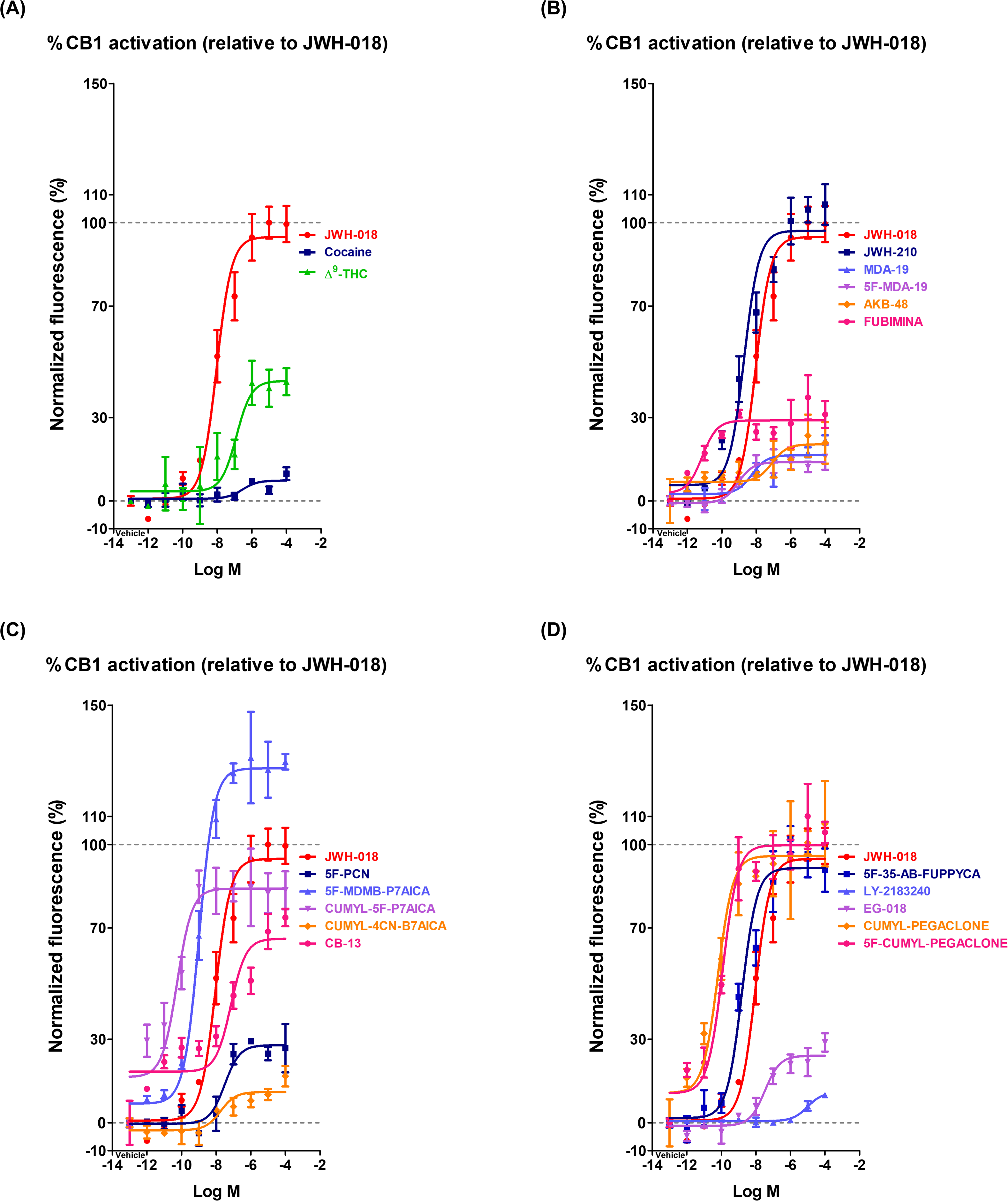
Dose-response curves of CB1-mediated Ca^2+^ signaling induced by SCRAs *in vitro*. A) Validation of the assay system using JWH-018 (full agonist), A^9^-THC (partial agonist), and cocaine (negative control). Intracellular Ca^2+^ responses were measured in CB1-Ga15 stably expressing Chem-1 cells and expressed as % △F/F_o_. JWH-018 and A9-THC induced dose-dependent increases in Ca^2+^ signaling, whereas cocaine did not elicit a measurable response. B–D) Dose-response curves of intracellular Ca^2+^ signaling induced by 16 SCRAs (0.001 nM-100 pM). Basal fluorescence was recorded for 60 s, followed by SCRA addition and measurement for an additional 60 s. Responses were normalized to the maximal effect of JWH-018 (100%). Dose-response curves were fitted using a four-parameter logistic nonlinear regression model to determine EC_50_ and *E*_max_ values.

Of the 16 SCRAs tested, dose-response curves were established for 11 SCRAs: JWH-018, JWH-210, 5F-PCN, 5F-MDMB-P7AICA, CUMYL-5F-P7AICA, CB-13, 5F-3,5-AB-PFUPPYCA, LY-2183240, EG-018, CUMYL-PEGACLONE, and 5F-CUMYL-PEGACLONE (Figs. 4B-D and S1; Table 1). The remaining five compounds, MDA-19, 5F-MDA-19, AKB-48, FUBIMINA, and CUMYL-4CN-B7AICA, showed weak CB1 agonistic responses, precluding reliable dose-response curve fitting.

**Table 1.** Summary of *in vitro* pharmacological parameters of SCRAs in the CB1-mediated Ca^2+^ assay. EC_50_ and *E*_max_ values were determined from dose-response curves obtained in CB1-Ga15 stably expressing Chem-1 cells following SCRA treatment (0.001 nM-100 μM). Intracellular Ca^2+^ responses were normalized to the maximal effect of JWH-018 (100%). Dose-response curves were fitted using a four-parameter logistic nonlinear regression model. SCRAs with poor curve fitting (R^2^ < 0.7) were excluded from analysis and designated as not determined (ND).

| No. | SCRAs | EC<sub>50</sub> (95% CI) | E<sub>max</sub> |
| --- | --- | --- | --- |
| 1 | JWH-018 | 11.67 nM (6.77-20.08 nM) | 100% |
| 2 | JWH-210 | 2.84 nM (1.63-4.98 nM) | 107% |
| 3 | MDA-19 | ND | ND |
| 4 | 5F-MDA-19 | ND | ND |
| 5 | AKB-48 | ND | ND |
| 6 | FUBIMINA | ND | ND |
| 7 | 5F-PCN | 27.86 nM (8.02-96.97 nM) | 29% |
| 8 | 5F-MDMB-P7AICA | 0.90 nM (0.55-1.45 nM) | 131% |
| 9 | CUMYL-5F-P7AICA | 0.0022 nM (0.00037-0.0032 nM) | 85% |
| 10 | CUMYL-4CN-B7AICA | ND | ND |
| 11 | CB-13 | 5.40 nM (1.76-16.53 nM) | 74% |
| 12 | 5F-3,5-AB-PFUPPYCA | 1.58 nM (0.79-3.19 nM) | 102% |
| 13 | LY-2183240 | 6792.04 nM (3548.13-13001.70 nM) | 10% |
| 14 | EG-018 | 115.61 nM (36.78-363.92 nM) | 29% |
| 15 | CUMYL-PEGACLONE | 0.074 nM (0.022-0.024 nM) | 107% |
| 16 | 5F-CUMYL-PEGACLONE | 0.087 nM (0.041-0.18 nM) | 110% |

### Several SCRAs produced dose-dependent catalepsy in mice

Separate groups of mice received each SCRA, and catalepsy was assessed to evaluate CB1 agonistic effects (Fig. 5). Among Class 1 compounds, the reference full agonist JWH-018 produced marked catalepsy from 0.5 mg/kg (Fig. 5A), and the structurally similar JWH-210 produced strong responses over a comparable dose range (Fig. 5B). Class 1B compounds MDA-19 and 5F-MDA-19 produced no significant catalepsy under the tested conditions (Figs. 5C and D).

**Fig. 5.**
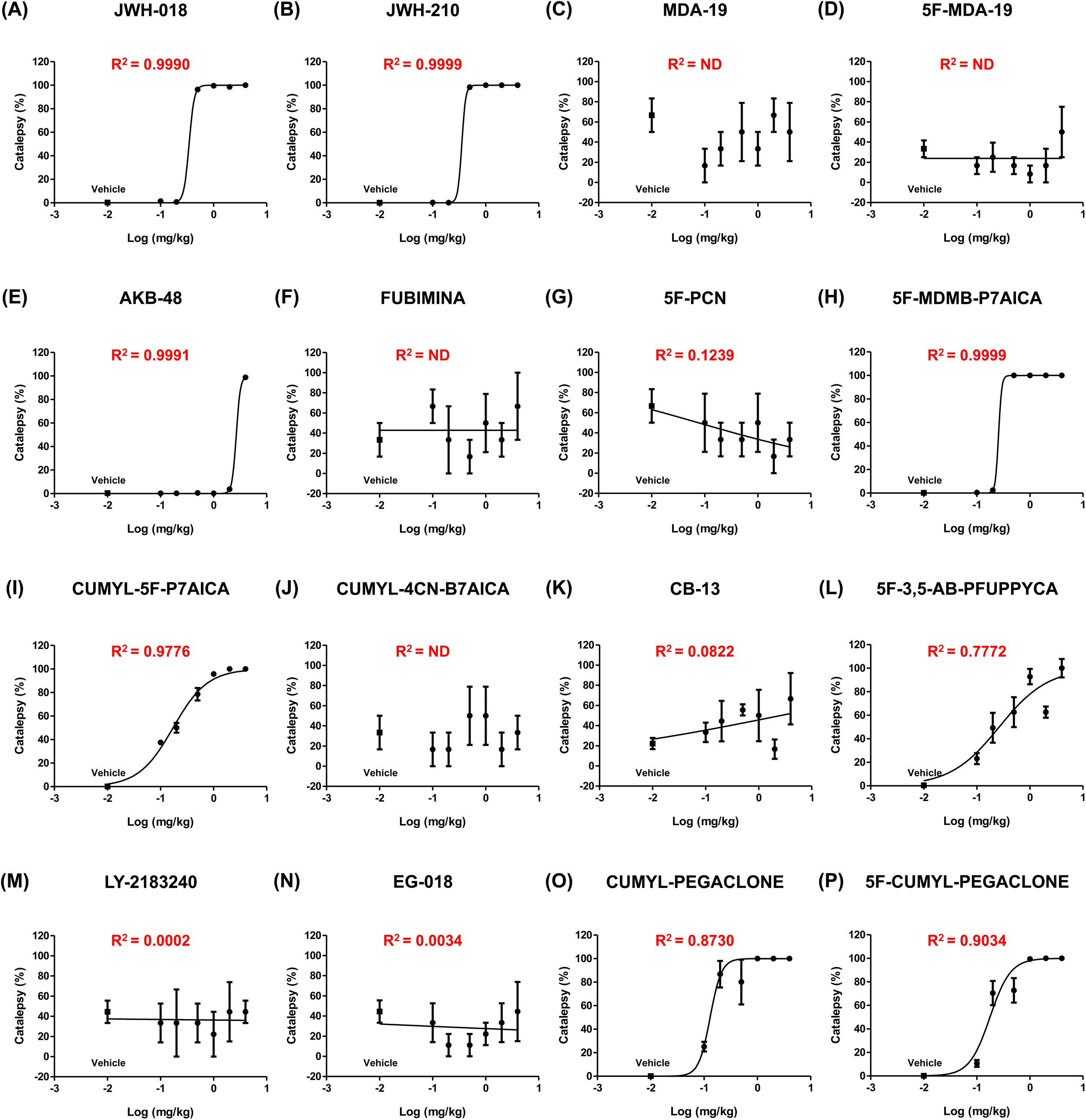
Dose-response curves of cataleptic responses induced by individual SCRAs. A–P) Cataleptic responses were measured by the bar test at 90 min after administration of each compound (0.1–4 mg/kg). The duration of catalepsy was recorded with a maximum cut-off time of 300 s. Responses were normalized to the maximal response of JWH-018, which was defined as 100%. Dose-response curves were fitted using a four-parameter logistic nonlinear regression model to calculate ED_50_ and *E*_max_ values. SCRAs with poor curve fitting (R^2^ < 0.7) were excluded from analysis. Data are presented as the mean ± SEM (*n* = 3).

AKB-48 (Class 2) produces marked catalepsy only at 4 mg/kg, the highest dose examined, while FUBIMINA (Class 3) and 5F-PCN (Class 4) showed no significant responses (Figs. 5E–G). Within Class 5, 5F-MDMB-P7AICA produced strong catalepsy from 0.5 mg/kg, and CUMYL-5F-P7AICA produced catalepsy from the lowest dose tested, 0.1 mg/kg, with duration increasing dose-dependently (Figs. 5H and I). CUMYL-4CN-B7AICA produced no significant effect (Fig. 5J).

CB-13 (Class 6), 5F-3,5-AB-PFUPPYCA (Class 7), LY-2183240 (Class 8), and EG-018 (Class 9) produced weak or no significant catalepsy. However, 5F-3,5-AB-PFUPPYCA produced measurable effects with a markedly lower maximal response (Figs. 5K–N; Table 2). The Class 10 compounds CUMYL-PEGACLONE and 5F-CUMYL-PEGACLONE produced clear dose-dependent catalepsy from the lowest administered dose (Figs. 5O and P).

**Table 2.**
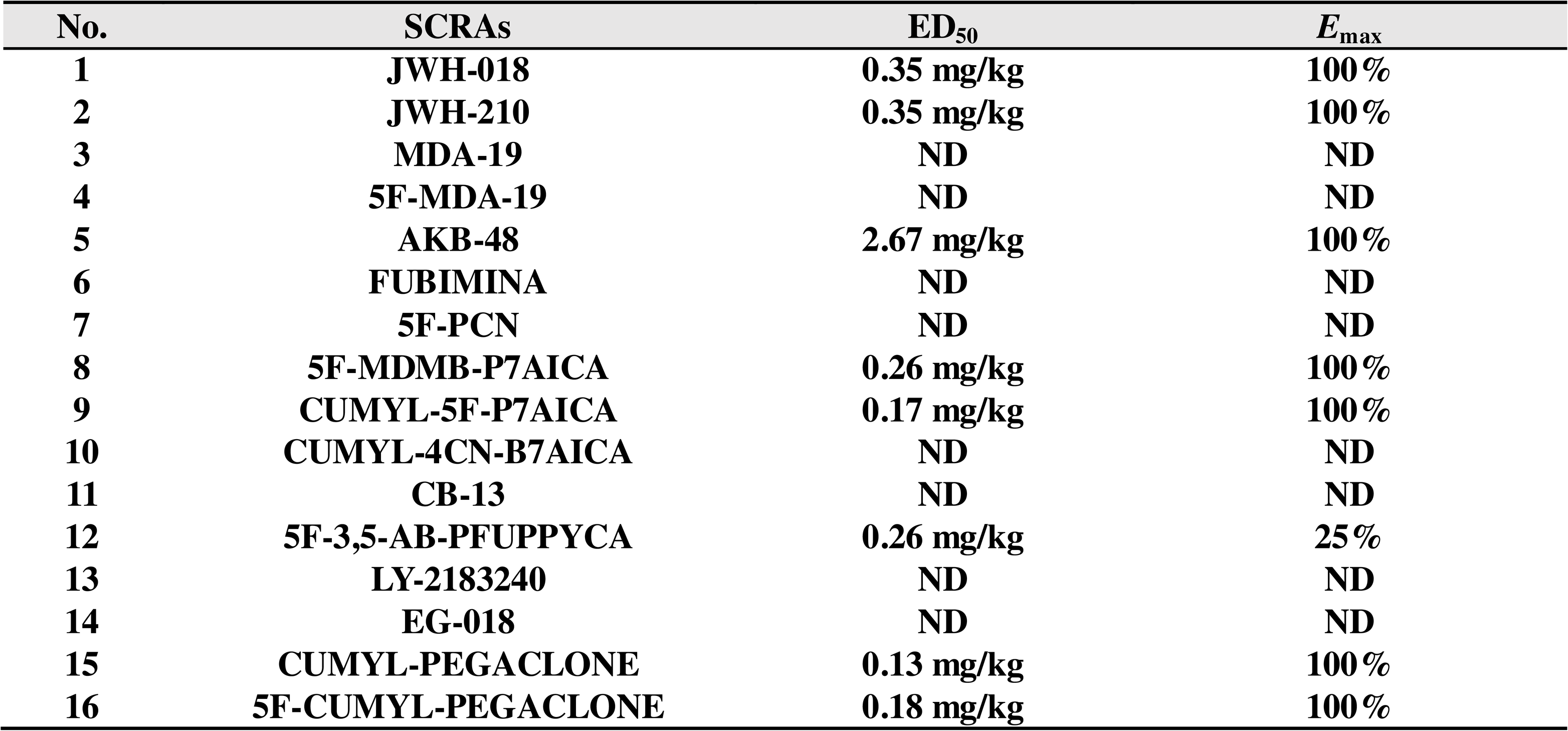
Summary of *in vivo* pharmacological parameters of SCRAs in the catalepsy test. ED_50_ and *E*_max_ values were determined from dose-response curves obtained in the bar test performed 90 min after drug administration (0.1–4 mg/kg). Cataleptic responses were normalized to the maximal effect of JWH-018 (100%). Dose-response curves were fitted using a four-parameter logistic nonlinear regression model. SCRAs with poor curve fitting (R^2^ < 0.7) were excluded from analysis and designated as not determined (ND).

Dose-response curves were established for eight SCRAs: JWH-018, JWH-210, AKB-48, 5F-MDMB-P7AICA, CUMYL-5F-P7AICA, 5F-3,5-AB-PFUPPYCA, CUMYL-PEGACLONE, and 5F-CUMYL-PEGACLONE (Fig. 5; Table 2). The remaining eight compounds, MDA-19, 5F-MDA-19, FUBIMINA, 5F-PCN, CUMYL-4CN-B7AICA, CB-13, LY-2183240, and EG-018, produced no quantifiable cataleptic responses, precluding reliable curve fitting.

### *In vitro*-*in vivo* pharmacological comparison

Correlation analyses related *in vivo* ED_50_ values from the catalepsy test to *in vitro* EC_50_ and *E*_max_ values. Spearman analysis showed a significant correlation between *in vivo* ED_50_ and *in vitro* EC_50_ values (Fig. 6A), whereas Pearson analysis of the log-transformed values showed no significant association (Fig. 6B). No significant correlation was found between *in vivo* ED_50_ and *in vitro E*_max_ (Fig. 6C).

**Fig. 6.**
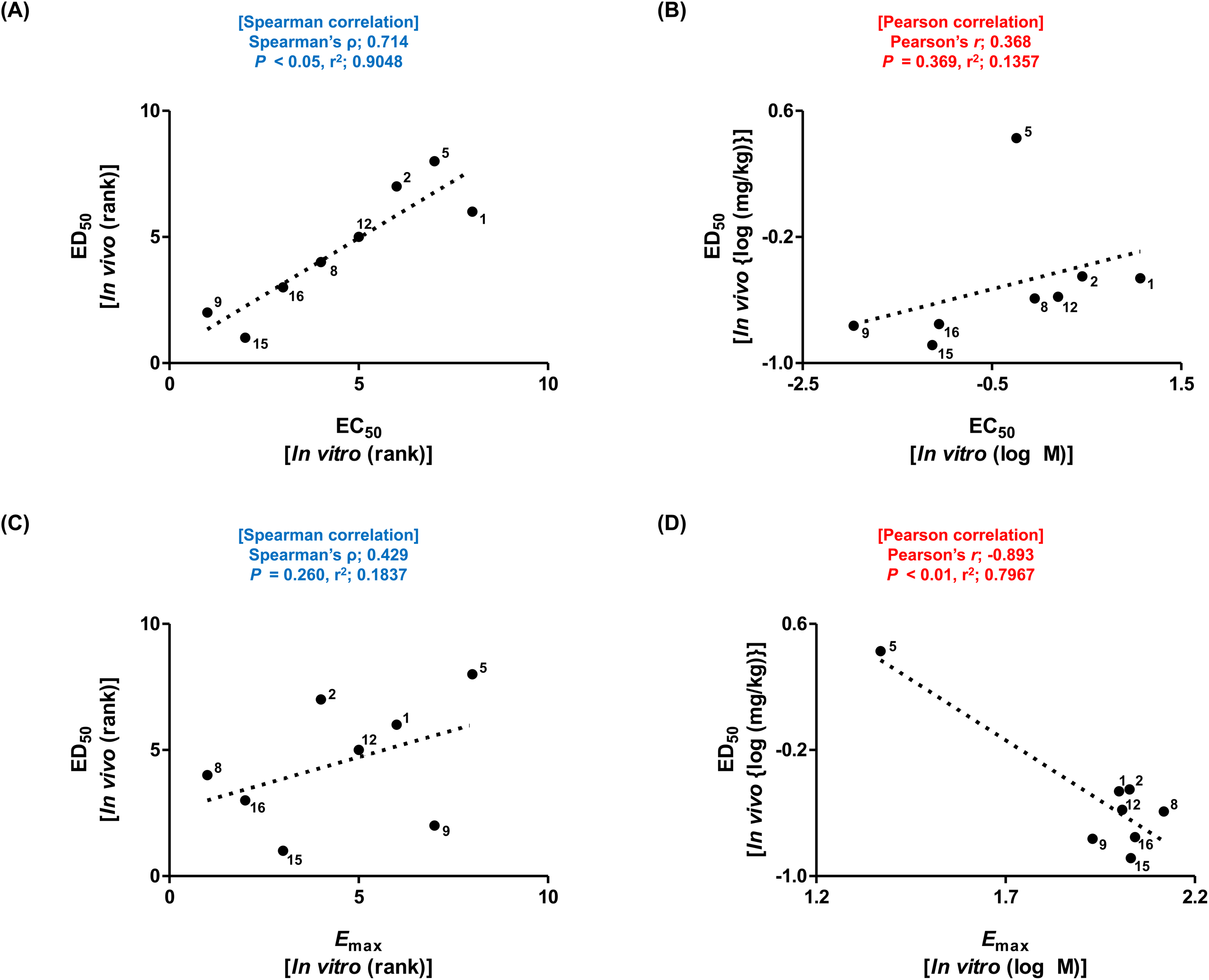
Correlation analysis between *in vitro* and *in vivo* pharmacological parameters of SCRAs. A) Correlation between *in vitro* EC_50_ and *in vivo* ED_50_ values analyzed using Spearman rank correlation. A strong positive correlation was observed, indicating that SCRAs with higher potency *in vitro* tend to exhibit greater potency *in vivo*. B) Correlation between log-transformed *in vitro* EC_50_ and *in vivo* ED_50_ values analyzed using Pearson correlation. No significant linear relationship was observed. C) Correlation between *in vitro E*_max_ and *in vivo* ED_50_ values analyzed using Spearman rank correlation. No significant association was observed. D) Correlation between log-transformed *in vitro E*_max_ and *in vivo* ED_50_ values analyzed using Pearson correlation. Although a linear trend was observed, the relationship was largely influenced by a single data point, suggesting limited reliability. Each data point represents an individual SCRA. Only SCRAs with determined values (non-ND) were included in the analysis. SCRA numbering is as follows: 1, JWH-018; 2, JWH-210; 5, AKB-48; 8, 5F-MDMB-P7AICA; 9, CUMYL-5F-P7AICA; 12, 5F-3,5-AB-PFUPPYCA; 15, CUMYL-PEGACLONE; 16, 5F-CUMYL-PEGACLONE.

Analysis of the log-transformed *in vivo* ED_50_ and *in vitro E*_max_values suggested a possible association, but this was driven largely by a single data point and was considered unreliable (Fig. 6D).

### Molecular docking identified different CB1 residue interaction profiles among SCRAs

Docking against the active-state CB1 structure (PDB ID: 6N4B) was used to examine ligand interactions with HIS178, ARG182, PHE189, PHE200, PHE268, TRP279, and TRP356 (Fig. 7A). PHE200 and TRP356 lie at the CB1 toggle switch, PHE268 and TRP279 contribute to ligand stabilization, and HIS178, PHE189, and ARG182 lie within or near the ligand-binding pocket. Docking poses exhibited varying combinations of tt-tt and other residue interactions, with distinct profiles observed among SCRAs differing in *in vitro* potency and efficacy. Quantitative analysis included the 11 SCRAs with determinable ECdd and *E_max_* values (Fig. 7B-L). Each ligand-residue interaction was categorized as no interaction, other interaction, or tt-tt interaction for clustering and PCA.

**Fig. 7.**
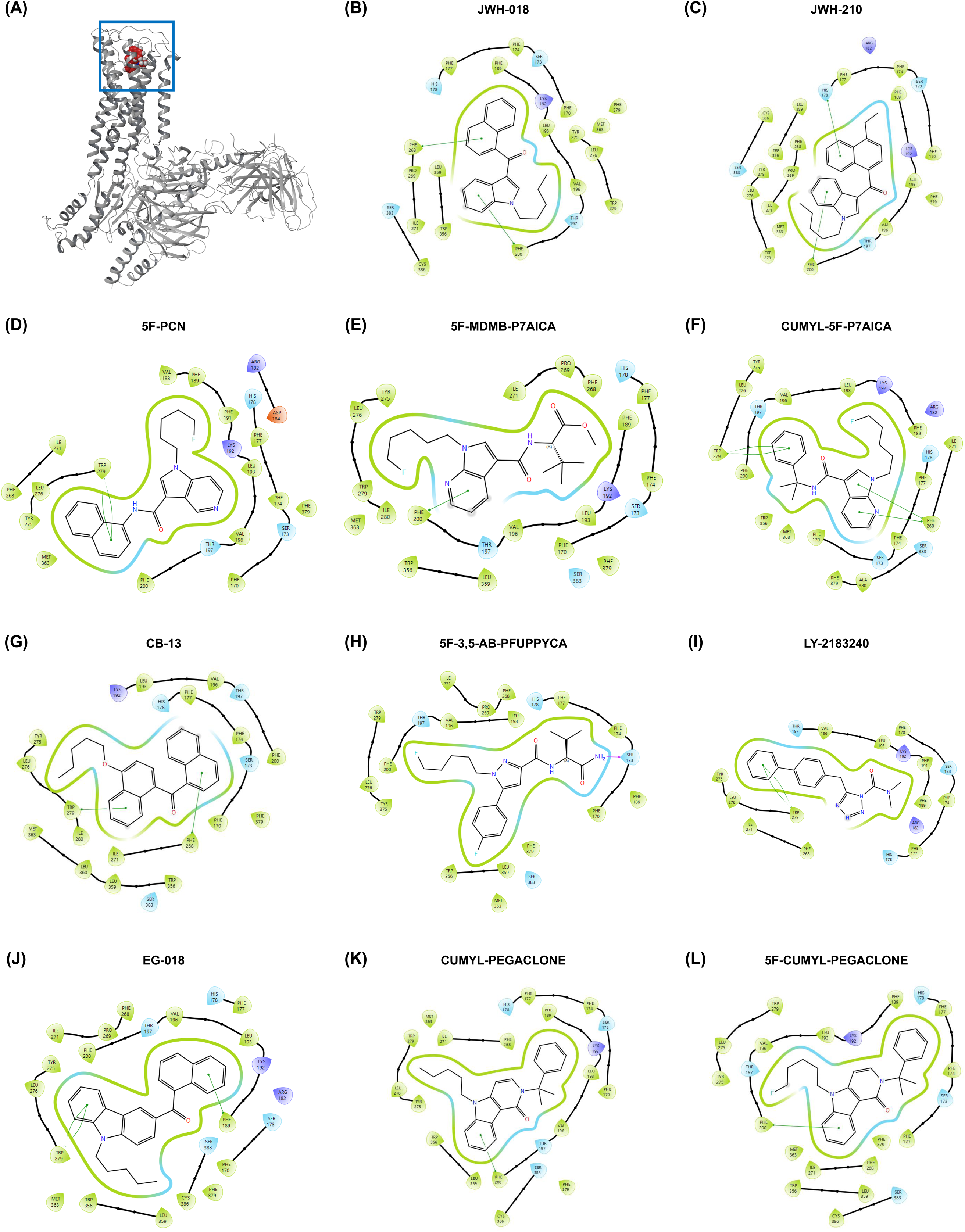
Molecular docking and ligand-residue interaction profiles of SCRAs at the CB1 receptor. Molecular docking was performed using the active-state CB1 receptor coupled to the heterotrimeric G_i_ protein (PDB; 6N4B). A) Overall structure of the CB1-G_i_ complex, with the CB1 receptor shown in gray. The orthosteric ligand-binding pocket is indicated by the blue box, and JWH-018, used as the representative ligand, is shown in red. B–L) Two-dimensional ligand-residue interaction diagrams of the 11 SCRAs for which reliable *in vitro* EC_50_ and *E*_max_ values were determined: (B) JWH-018; (C) JWH-210; (D) 5F-PCN; (E) 5F-MDMB-P7AICA; (F) CUMYL-5F-P7AICA; (G) CB-13; (H) 5F-3,5-AB-PFUPPYCA; (I) LY-2183240; (J) EG-018; (K) CUMYL-PEGACLONE; and (L) 5F-CUMYL-PEGACLONE.

### CB1 residue interaction profiles were associated with *in vitro* efficacy

Hierarchical clustering of the ligand–residue interaction profiles divided the 11 SCRAs into three groups (Fig. 8A). The first cluster (red) included JWH-018, JWH-210, 5F-MDMB-P7AICA, 5F-3,5-AB-PFUPPYCA, CUMYL-PEGACLONE, and 5F-CUMYL-PEGACLONE, which showed high *E*_max_. CUMYL-5F-P7AICA and CB-13 formed the second cluster (green), characterized by intermediate *E*_max_, whereas 5F-PCN, LY-2183240, and EG-018 formed the third cluster (purple), characterized by the lowest *E*_max_.

**Fig. 8.**
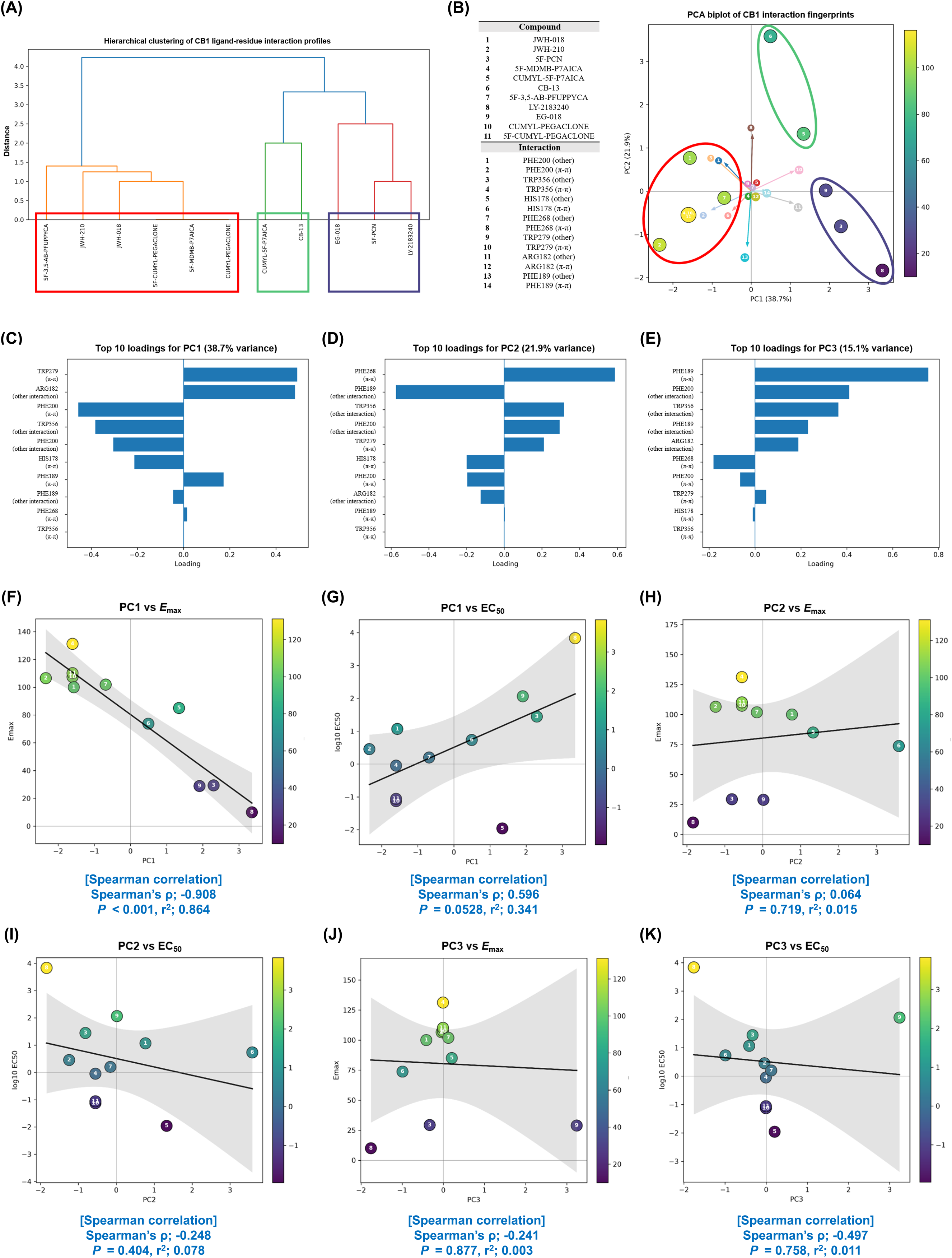
Hierarchical clustering and principal component analysis of CB1 ligand-residue interaction profiles and their relationships with *in vitro* pharmacological activity. Residue-level interaction fingerprints were generated for 11 SCRAs based on their interactions with seven key residues within the CB1 ligand-binding pocket (HIS178, ARG182, PHE189, PHE200, PHE268, TRP279, TRP356). A) Hierarchical clustering of CB1 ligand-residue interaction profiles. The 11 SCRAs were separated into three major clusters, indicated by red, green, and purple boxes. B) PCA biplot of CB1 ligand-residue interaction fingerprints showing the distribution of individual SCRAs according to PC1 and PC2 and the direction of interaction-feature loadings. Compounds are colored according to their *in vitro E*_max_ values. C– E) Top 10 feature loadings contributing to PC1, PC2, and PC3, respectively. F–K) Relationships between principal component scores and *in vitro* pharmacological parameters: (F) PC1 versus *E*_max_, (G) PC1 versus log_10_ EC_50_, (H) PC2 versus *E*_max_, (I) PC2 versus log_10_ EC_50_, (J) PC3 versus *E*_max_, and (K) PC3 versus log_10_ EC_50_. Compounds in panels F–K are colored according to their *in vitro E*_max_ values. Solid lines represent fitted regression lines, and gray-shaded areas indicate the 95% confidence intervals. Associations between principal component scores and pharmacological parameters were evaluated using Spearman’s rank correlation analysis, followed by Benjamini-Hochberg false discovery rate (FDR) correction for multiple comparisons.

PCA of the same profiles showed that PC1, PC2, and PC3 explained 38.7%, 21.9%, and 15.1% of the total variance, respectively, accounting for 75.7% combined (Figs. 8B–E). The main PC1 loadings involved TRP279, ARG182, PHE200, TRP356, and HIS178 (Fig. 8C).

A strong association was observed between PC1 and *in vitro E*_max_, which remained significant following Benjamini-Hochberg FDR adjustment (FDR-adjusted *q* < 0.05; Fig. 8F). PC1 was not correlated with logan ECll, and the association remained non-significant after FDR correction (Fig. 8G). No association of PC2 or PC3 with logo □ ECl l or *E*_max_ remained significant after correction (FDR-adjusted *q* > 0.05; Fig. 8H–K). Thus, PC1-*E*_max_ was the only association that remained significant after correction for multiple comparisons.

## Discussion and Conclusions

We investigated CB1 agonistic activity across 16 SCRAs using *in vitro* and *in vivo* approaches and integrated structural classification, molecular descriptor analysis, molecular docking, and residue-level interaction analysis to examine structure-activity relationships underlying differences in CB1 pharmacology. The 16 SCRAs were selected for their regulatory relevance across jurisdictions with different scheduling systems, including the United States, Australia, Japan, and the Republic of Korea [6–8,21–26]. These differences reflect the global challenge of regulating SCRAs because structurally similar compounds can differ markedly in pharmacology. Systematic evaluation of functional activity is therefore essential to complement structure-based regulatory approaches.

The SCRAs were classified by core scaffold and by linker, head, tail, and additional residue groups. Molecular fingerprint-based hierarchical clustering and similarity heatmap analysis supported this scheme, with structurally similar compounds clustering together [16], providing a structural basis for interpreting pharmacological differences among SCRAs.

The Ca^2+^-based *in vitro* assay uses Ga15-mediated coupling to convert GPCR activation into measurable intracellular Ca^2+^ signaling, providing a sensitive, high-throughput platform for functional analysis [20,28]. Assay validation with the full agonist JWH-018, partial agonists A9-THC, and negative control cocaine demonstrated clear differentiation of their pharmacological profiles. The EC_50_ and *E*_max_ values were consistent with published values for well-characterized SCRAs, particularly JWH analogues [29,30]. The individual concentration-response curves in the supplementary figures show the wide range of CB1 responses across the tested compounds, from highly efficacious agonists to compounds for which reliable EC_50_ and *E*_max_ values could not be determined. MDMB- and CUMYL-like SCRAs were highly potent and efficacious. Two discrepancies stand out: CUMYL-4CN-B7AICA showed no measurable CB1 agonism despite sharing a structural class with highly active compounds, and the dose-response curve for AKB-48 fitted poorly, preventing accurate parameter estimation. Such differences may reflect pathway-specific signaling, as some SCRAs show functional selectivity in signaling through G proteins versus β-arrestins [29,30].

*In vivo*, JWH-like compounds produced strong catalepsy consistent with their known full agonism at CB1 [31], and 5F-MDMB-P7AICA and CUMYL-like compounds were likewise highly potent and efficacious [32]. AKB-48 required relatively high doses for maximal catalepsy, consistent with reports of lower potency but retained full efficacy [29,30], implying that greater systemic exposure is needed for sufficient receptor occupancy. 5F-3,5-AB-PFUPPYCA showed moderate potency but low maximal efficacy *in vivo*, suggesting partial agonism or signaling bias, as reported for structurally related SCRAs [29]; this compound is also frequently conflated with 5F-AB-FUPPYCA in the literature, making comparison with earlier work difficult [33]. MDA-19, 5F-MDA-19, FUBIMINA, and CUMYL-4CN-B7AICA produced no measurable catalepsy, which may indicate weak or absent CB1 agonism or activity at alternative targets [7,8,34]. High-dose CB1 agonists can induce aversive responses that may mask catalepsy [35,36]. The dose range was therefore selected to balance sensitivity against non-specific or aversive. Curves with poor fitting (R^2^ < 0.7) were excluded from quantitative analysis because they were considered unreliable for nonlinear parameter estimation [37].

Despite broadly similar *in vitro* and *in vivo* patterns, CB-13 and EG-018 displayed *in vitro* activity without detectable cataleptic responses. This discrepancy likely reflects the inability of the engineered *in vitro* system to reproduce *in vivo* pharmacology fully, including pharmacokinetic factors such as blood-brain barrier penetration, systemic exposure, distribution, and metabolic stability, together with differences in receptor coupling and signal amplification between the CB1-Ga15 assay and native G_i_/o-mediated CB1 signaling [38,39]. Receptor-level activity therefore need not translate into behavioral effects when compound disposition and signaling environment differ from the cell-based system.

Spearman analysis showed a strong positive association between *in vitro* EC_50_ and *in vivo* ED_50_, indicating that receptor-level potency may partly reflect potency in the catalepsy test. The absence of a significant association between *in vitro E*_max_ and *in vivo* ED_50_ supports the distinction between potency, governed by ligand affinity and receptor-effector coupling, and efficacy, which is influenced by receptor density, signal amplification, and coupling efficiency [40]. Pearson analysis of log-transformed data was not significant, probably because of outliers, particularly AKB-48, and the apparent *E*_max_-ED_50_ correlation was driven by a single influential point, a well-described source of misleading regressions [41,42]. The in *vitro*-*in vivo* potency relationship should therefore be read as support for the pharmacological relevance of the assay rather than as evidence that receptor-level activity alone determines behavioral potency.

Descriptor-based analysis gave a more quantitative view of these relationships. Potency-related associations did not survive FDR correction, whereas several efficacy-related ones did: the order-4 charge index remained associated with *in vitro E*_max_, and aromaticity-, molecular weight-, and hydrogen-related descriptors with *in vivo E*_max_. Within this dataset, therefore, charge distribution, aromaticity, and molecular size appear more closely linked to maximal response than to potency.

As mathematical measures of structural and physicochemical properties, molecular descriptors do not directly define mechanisms but may represent broader features affecting ligand orientation, receptor stabilization, and steric and hydrophobic complementarity within the ligand-receptor complex. This is consistent with quantitative structure-activity relationship principles emphasizing the combined importance of structural and electronic properties in ligand-receptor interactions [15,43,44].

Given the small number of SCRAs relative to the descriptors examined, the observed associations warrant consideration as exploratory rather than definitive. Confirmation using larger SCRA datasets with distinct structural compositions is needed to determine whether the efficacy-related associations generalize.

Docking and residue-level interaction analyses using the active-state CB1-G_i_ structure addressed the receptor-level basis of these differences. Structural studies of agonist-bound CB1 show that PHE200^3.36^ and TRP356^6.48^ form an activation-associated “toggle switch” that undergoes coordinated conformational rearrangement during receptor activation, and that the potent SCRA MDMB-Fubinaca engages these residues to stabilize an active conformation associated with outward TM6 movement and G-protein engagement [18]. MDMB-Fubinaca also forms aromatic interactions with TRP279^5.43^, supporting an aromatic network of PHE200, TRP279, and TRP356 in active-state stabilization [18].

Consistent with this framework, hierarchical clustering of the ligand–residue interaction profiles broadly separated SCRAs by *in vitro* efficacy: the first cluster (JWH-018, JWH-210, 5F-MDMB-P7AICA, 5F-3,5-AB-PFUPPYCA, CUMYL-PEGACLONE, and 5F-CUMYL-PEGACLONE) had relatively high *E*_max_, whereas the third (5F-PCN, LY-2183240, and EG-018) had substantially lower maximal responses. Clustering alone does not establish a statistical relationship, but it suggests that compounds with similar predicted interaction profiles may share functional characteristics.

PCA supported this relationship. PC1, which explained 38.7% of the variance and was strongly influenced by interactions involving TRP279, ARG182, PHE200, TRP356, and HIS178, was strongly associated with *in vitro E*_max_ and was the only principal component-parameter association to survive Benjamini-Hochberg FDR correction; PC1 was only non-significantly related to log_10_ EC_50_, and PC2 and PC3 were unrelated to either parameter. The major variation captured by the residue interaction fingerprints therefore tracked efficacy more closely than potency.

This is consistent with structural evidence that ligand interactions within the CB1 binding pocket influence signaling efficacy. Beyond the PHE200-TRP356 toggle switch, structural, molecular dynamics, and signaling studies indicate that interactions with surrounding regions propagate conformational change toward the intracellular surface and affect G protein activation [45]. The prominent contributions of PHE200, TRP356, and TRP279 to PC1 are therefore mechanistically compatible with established CB1 activation mechanisms [18,45].

The analysis does not, however, show that any individual residue independently determines efficacy. The strong PC1-*E*_max_ relationship instead suggests that the collective interaction pattern across activation- and ligand-stabilizing residues distinguishes high-from low-efficacy SCRAs better than any single contact.

LY-2183240 is an important exception. Its *in vivo* cataleptic response was ND, and although a concentration-response curve was obtained in the CB1-Ga15 assay, the response was low and comparable to the negative control cocaine. LY-2183240 is primarily an inhibitor of endocannabinoid metabolism, potently inhibiting FAAH and other brain serine hydrolases, rather than a direct CB1 agonist [46]; the weak response seen here is therefore insufficient to classify it as a functional SCRA.

The descriptor- and interaction-based analyses therefore converge on efficacy as the more informative pharmacological dimension in this dataset: FDR-robust associations linked molecular descriptors to both *in vitro* and *in vivo E*_max_, and residue-interaction PCA independently identified a pattern significantly associated with *in vitro E*_max_. Despite operating at different levels of structural information, both point to maximal CB1 response rather than potency as the property most closely tied to measurable molecular features.

Several limitations apply. The number of descriptors was large relative to the compound set; thus, some descriptor-level associations may be dataset-specific despite FDR adjustment. Docking yields predicted rather than experimentally resolved binding modes, and the functional contribution of individual residues was not validated by site-directed mutagenesis or other residue-specific functional experiments. The CB1-Ga15 Ca^2+^ assay is an engineered signaling system that may not reproduce native G_i/o_ coupling or other CB1 pathways. Additionally, the lack of pharmacokinetic measurements limited mechanistic interpretation of the discordant responses observed across experimental systems.

The associations reported here should not yet be considered predictive of efficacy. Validation requires larger and more diverse datasets, native CB1 assays, mutagenesis, simulations, and pharmacokinetics. Nevertheless, our findings effectively demonstrate substantial differences in CB1 potency and efficacy among SCRAs, with efficacy more closely tied to molecular and receptor-interaction features. Integrating functional pharmacology with structural and receptor-interaction analyses may help prioritize emerging compounds for further pharmacological and behavioral evaluation.

## Supporting information

Supplemental Figure 1-2, additional method, and results

## Author Contributions

Hyeokjun Kwon; Data Curation, Formal analysis, Investigation, Methodology, Validation, Visualization, Writing – Original Draft. Chul kyu Lee; Data Curation, Investigation, Validation. Jae-Hwan Kwak; Investigation, Methodology, Supervision, Writing – Review & Editing. Jaesuk Yun; Conceptualization, Funding acquisition, Project administration, Resources, Software, Supervision, Writing – Review & Editing.

## Acknowledgements

This work was supported by the Ministry of Food and Drug Safety (23212MFDS217), the Bio&Medical Technology Development Program of the National Research Foundation (NRF) funded by the Korean government (MSIT) (No. RS-2024-00440787), the NRF of Korea grant funded by the Korean government (MSIT) (No. RS-2025-02273102), the Basic Science Research Program through the NRF funded by the Ministry of Education (RS-2024-00460411).

## Data Availability Statement

The data that support the findings of this study are available from the corresponding author upon reasonable request. Some data may not be available because of privacy or ethical restrictions.

