## Supplemental Figure 1-2, additional method, and results for "Integrated pharmacological and structural profiling reveals CB1 residue interaction patterns associated with synthetic cannabinoid receptor agonist efficacy"

### Supporting Information

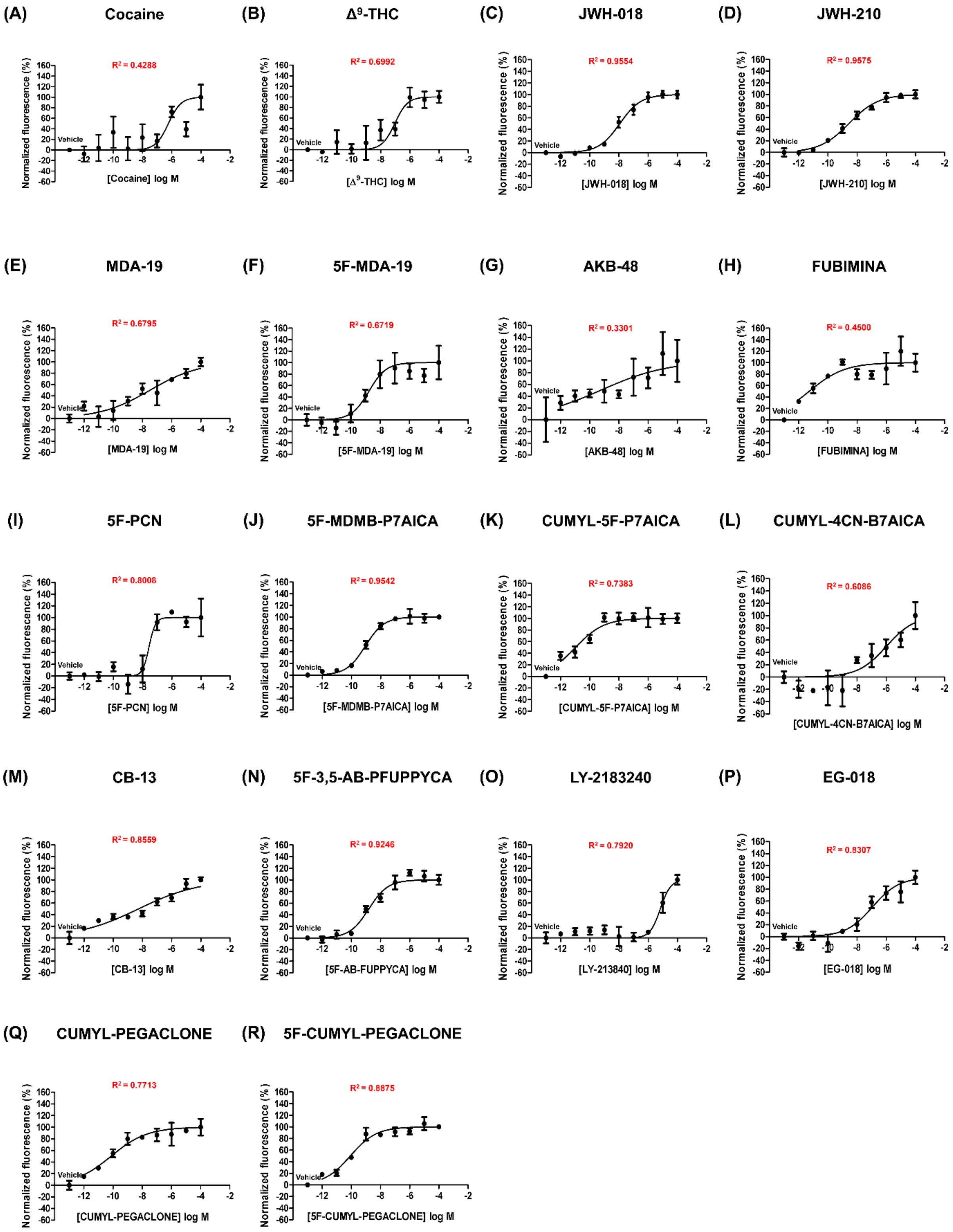


**Fig. S1. Dose-response curves of SCRAs in the CB1-mediated $Ca^{2+}$ assay.** A-B) Dose-response curves for cocaine (negative control) and $\Delta^9$-THC (partial agonist), validating assay specificity and dynamic range. C-R) Dose-response curves for 16 SCRAs. Data are presented as the mean ± SEM (n = 3).

**Descriptor-based correlation analysis**

To investigate structural determinants of CB1 agonism, molecular descriptors representing physicochemical and topological properties were calculated using the Mold2 descriptor set and correlated with four pharmacological outcomes: *in vitro* $EC_{50}$, *in vitro* $E_{max}$, *in vivo* $ED_{50}$, and *in vivo* $E_{max}$. Spearman's rank correlation was used to capture potential non-linear relationships. Results were visualized as heatmaps comparing the relative contributions of the descriptors to pharmacological activity.

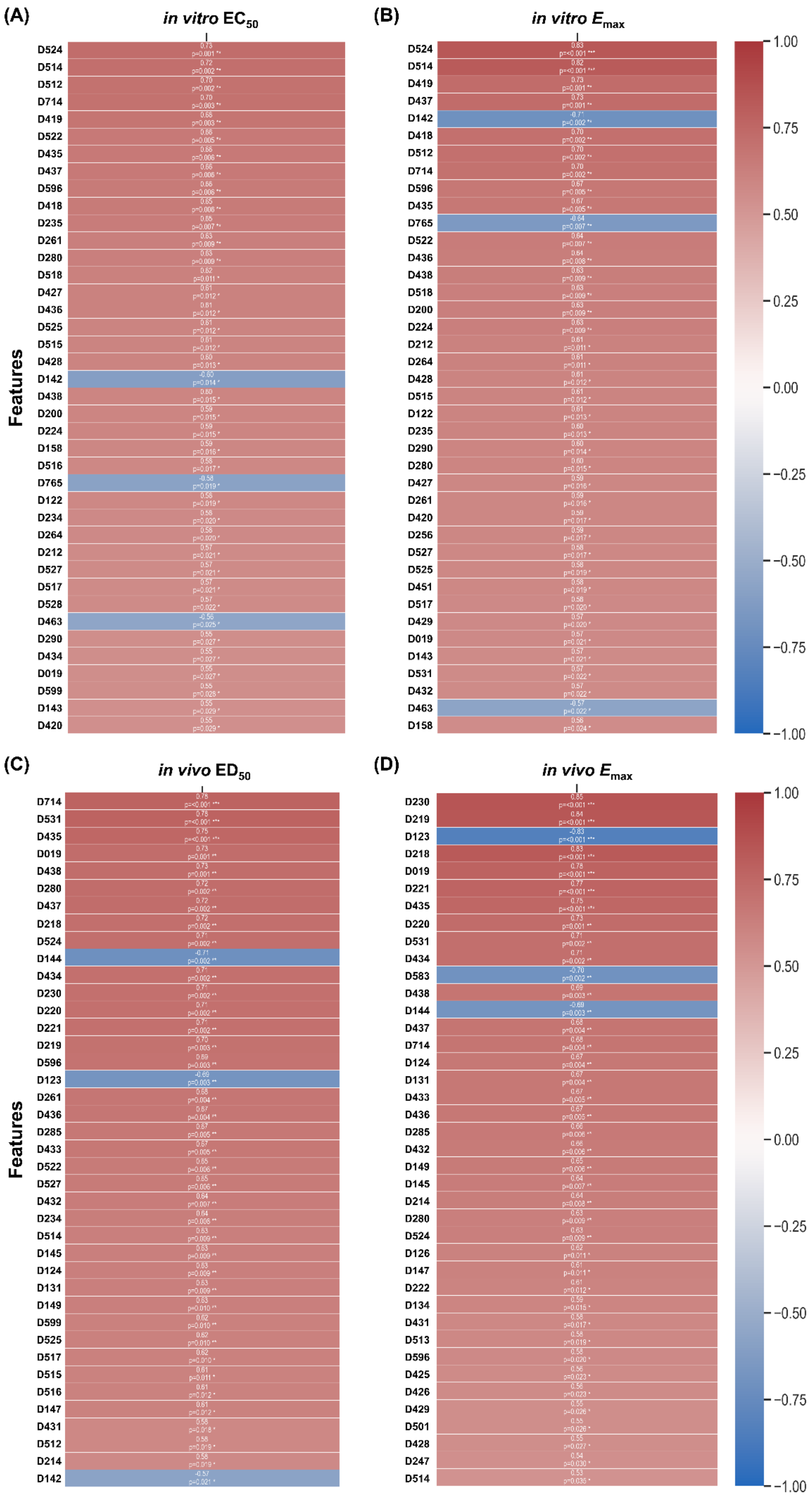
(A)
*in vitro* $EC_{50}$
Features
D524
D514
D512
D714
D419
D522
D435
D437
D596
D418
D235
D261
D280
D518
D427
D436
D525
D515
D428
D142
D438
D200
D224
D158
D516
D765
D122
D234
D264
D212
D527
D517
D528
D463
D290
D434
D019
D599
D143
D420
(B)
*in vitro* $E_{max}$
D524
D514
D419
D437
D142
D418
D512
D714
D596
D435
D765
D522
D436
D438
D518
D200
D224
D212
D264
D428
D515
D122
D235
D290
D280
D427
D261
D420
D256
D527
D525
D451
D517
D429
D019
D143
D531
D432
D463
D158
1.00
0.75
0.50
0.25
0.00
−0.25
−0.50
−0.75
−1.00
(C)
*in vivo* $ED_{50}$
Features
D714
D531
D435
D019
D438
D280
D437
D218
D524
D144
D434
D230
D220
D221
D219
D596
D123
D261
D436
D285
D433
D522
D527
D432
D234
D514
D145
D124
D131
D149
D599
D525
D517
D515
D516
D147
D431
D512
D214
D142
(D)
*in vivo* $E_{max}$
D230
D219
D123
D218
D019
D221
D435
D220
D531
D434
D583
D438
D144
D437
D714
D124
D131
D433
D436
D285
D432
D149
D145
D214
D280
D524
D126
D147
D222
D134
D431
D513
D596
D425
D426
D429
D501
D428
D247
D514
1.00
0.75
0.50
0.25
0.00
−0.25
−0.50
−0.75
−1.00

**Fig. S2. Heatmap of Spearman correlation between molecular descriptors and pharmacological parameters of SCRAs.** A) *in vitro* $EC_{50}$, B) *in vitro* $E_{max}$, C) *in vivo* $ED_{50}$, D) *in vivo* $E_{max}$. Spearman correlation analysis was performed to evaluate the association between physicochemical descriptors and pharmacological activity. Molecular descriptors, including indices related to charge distribution, electronegativity, molecular topology, and atomic composition, were calculated for each SCRAs. For each panel, correlation coefficients between individual descriptors and the corresponding pharmacological parameters are shown. Color intensity represents the magnitude and direction of correlation, with red indicating positive correlations and green indicating negative correlations.

**Descriptor-based correlation analysis shows that distinct structural features are associated with each pharmacological parameter.**

To evaluate the relationships between molecular properties and CB1 receptor-mediated activity, correlation analyses were performed using molecular descriptors (Fig. S2). To account for multiple comparisons, the resulting $P$ values were adjusted using the Benjamini-Hochberg false discovery rate (FDR) procedure, with an FDR-adjusted $q < 0.05$ considered statistically significant.

Descriptor-based analysis identified multiple physicochemical properties showing associations with pharmacological activity (Fig. S2). For *in vitro* $EC_{50}$, charge index descriptors (order-4 and order-2) showed the strongest associations, followed by the number of CH3R and CH4 groups; however, none of these associations remained statistically significant after FDR correction (FDR-adjusted $q > 0.05$). A similar pattern was observed for *in vitro* $E_{max}$, with charge index (order-4) showing the strongest associations, followed by atomic mass-related descriptors (length-4 and length-5), atomic Sanderson electronegativity indices (length-7), and topological descriptors such as the Balaban-type mean square vertex distance index. Among these descriptors, only charge index (order-4) remained significantly associated with *in vitro* $E_{max}$ after FDR correction (FDR-adjusted $q < 0.05$), whereas the other highly ranked descriptors did not retain statistical significance. In contrast, *in vivo* $ED_{50}$ was primarily associated with the number of CH3R and CH4 groups, followed by descriptors related to molecular charge distribution and electronegativity; however, none of these associations remained significant after FDR correction (FDR-adjusted $q >0.05$). For *in vivo* $E_{max}$, aromaticity-related descriptors, particularly the valence vertex connectivity index (order-1), showed the strongest correlations, followed by molecular weight- and hydrogen-related descriptors. Notably, several of these descriptors remained statistically significant after FDR correction (FDR-adjusted $q < 0.05$). Collectively, although several descriptor-level correlation patterns were observed across the four pharmacological parameters, statistically significant associations after FDR correction were restricted to efficacy-related parameters, including charge index (order-4) with *in vitro* $E_{max}$ and aromaticity-, molecular weight-, and hydrogen-related descriptors with *in vivo* $E_{max}$.
